# Haspin participates in Aurora phosphorylation at centromeres and contributes to chromosome congression in male mouse meiosis

**DOI:** 10.1101/2021.11.02.466959

**Authors:** I Berenguer, P López-Jiménez, I Mena, A Viera, J Page, C Maestre, M Malumbres, JA Suja, R Gómez

**Author notes:** These authors contributed equally to this work (I Berenguer, P López-Jiménez).

## Abstract

Chromosome segregation requires that centromeres properly attach to spindle microtubules. This is an essential step towards the accuracy of cell division and therefore must be precisely regulated in both mitosis and meiosis. One of the main centromeric regulatory signaling pathways is the Haspin-H3T3ph-chromosomal passenger complex (CPC) cascade, which is responsible for the recruitment of the CPC to the centromeres. In mitosis, Haspin kinase phosphorylates H3 at threonine 3 (H3T3ph), the essential histone mark that recruits the CPC whose catalytic component is Aurora B kinase. To date, no data has yet been presented about the action of the centromeric Haspin-H3T3ph-CPC pathway in mammalian male meiosis. We have analyzed the consequences of Haspin chemical inhibition in cultured spermatocytes using LDN-192960. Our *in vitro* studies suggest that Haspin kinase activity is required for proper chromosome congression during both meiotic divisions and for the recruitment of phosphorylated Aurora B at meiotic centromeres. These results have been confirmed by the characterization of the meiotic phenotype of the genetic mouse model *Haspin*^*-/-*^, which displays similar defects. In addition, our work demonstrates that the absence of H3T3ph histone mark does not alter SGO2 localization to meiotic centromeres. These results add new and relevant information regarding the regulation of centromere function during meiosis.

## INTRODUCTION

Chromosome instability and mis-segregation during mitosis and meiosis lead to the appearance of aneuploidies, a hallmark cause of tumorigenesis or prenatal death in vertebrates. The gain or loss of chromosomes often originates from dysregulation of centromere assembly, errors in kinetochore-microtubule interaction, or chromosome congression failures. These processes are highly regulated and well-orchestrated by several master regulators, such as PLK1 and Aurora B kinase (AURKB) among others, which perform their role in both mitosis (Lukasiewicz and Lingle, 2009; Nikonova et al., 2013; Schmucker and Sumara, 2014) and meiosis (Alfaro et al., 2021; Wellard et al., 2021). A recently described and poorly studied regulator is the kinase Haspin. Haspin (encoded by germ cell-specific gene 2: *Gsg2*) is an evolutionary conserved serine/threonine kinase (Cairo and Lacefield, 2020; Higgins, 2010) that was first described in mouse testis as a meiosis-specific gene, even though it is also expressed in somatic cells (Tanaka et al., 1999). Several substrates are known for this kinase, including histone H3 (through the phosphorylation at tyrosine 3, H3T3ph), histone macroH2A (through the phosphorylation at serine 137, macroH2AS137ph) and the centromeric protein CENP-T (phosphorylated at tyrosine 57, CENP-T T57ph) (Dai and Higgins, 2005; Maiolica et al., 2014; Markaki et al., 2009).

Most of the studies about Haspin have been conducted in somatic cells, where it was described as the main responsible regulator for the recruitment of the chromosomal passenger complex (CPC) to the inner centromere domain (ICD) through the phosphorylation of H3T3 (Higgins, 2010; Wang et al., 2010). The CPC is a multiprotein complex that possesses AURKB as the enzymatic subunit, playing an essential role during the regulation of chromosome segregation and cytokinesis (Krenn and Musacchio, 2015; van der Horst et al., 2015). The signaling network involving Haspin-mediated H3T3ph and Bub1-mediated histone H2A T120 phosphorylation (H2AT120ph) underlies the defined localization of AURKB at the mitotic ICD (Hadders et al., 2020; Liang et al., 2020). However, Haspin subcellular distribution pattern remains unclear. In somatic cells, Haspin coupled to GFP was detected at the centrosomes and in the nucleus, but its distribution seems diffuse (Dai et al., 2006; Dai et al., 2005). During interphase, Haspin presents a structural inactivation (Ghenoiu et al., 2013) and it is activated at prophase by two consecutive phosphorylations generated by CDK1 and PLK1 (Zhou et al., 2014). Once activated, Haspin phosphorylates H3T3 from prophase to anaphase over the entire chromatin, being more concentrated at the centromeres in metaphase (Dai and Higgins, 2005). H3T3ph then recruits the CPC to the ICD through the Survivin BIR domain (Kelly et al., 2010). Once in the centromere, AURKB generates a positive feedback loop and phosphorylates Haspin, promoting a higher accumulation of the CPC at centromeres (Wang et al., 2011). At the end of the mitotic metaphase, H3T3ph is dephosphorylated (Kelly et al., 2010) by the action of the PP1/Repo-Man complex and Haspin is inactivated (Qian et al., 2013; Vagnarelli et al., 2011).

Since its discovery, several publications have reported functional studies about Haspin in somatic cells. Haspin activity knockdown by RNAi induces the absence of H3T3ph and causes alterations in chromosome alignment (Dai et al., 2005) and premature loss of sister chromatid cohesion (Dai et al., 2009; Dai et al., 2005), suggesting a role of this kinase in centromeric cohesion and pointing to a potential link with Shugoshin 1 (SGO1). In this sense, several works colocalized H3T3ph with the cohesin subunit SA2 (Dai et al., 2006), and also suggested the association of Haspin with PDS5, a regulator of the sister chromatid cohesion (Carretero et al., 2013; Liang et al., 2018; Yamagishi et al., 2010). On the other hand, the chemical inhibition of this kinase in somatic animal and plant cells using different inhibitors such as 5-Iodotubercidin (5-Itu), CHR-6494 (CHR) and LDN-192960 (LDN) impairs AURKB localization to the centromeres (Cuny et al., 2012; Huertas et al., 2012; Patnaik et al., 2008; Wang et al., 2012). These data confirm the involvement of Haspin in the regulation of AURKB to fulfill chromosome segregation in mitosis (De Antoni et al., 2012; Kozgunova et al., 2016; Wang et al., 2012). Following this approach and the increasing development of chemical inhibitor drugs, strategies using Haspin as anti-cancer target are arising (Amoussou et al., 2018). Moreover, the up-regulation of this kinase in some cancer cells has corroborated a central role of Haspin in cell proliferation, increasing the interest of Haspin as a potential target for cancer treatment (Zhu et al., 2020).

While most studies on Haspin have focused on somatic cells, significantly fewer reports have been published about the distribution and functions of Haspin in meiosis (Cairo and Lacefield, 2020). Surprisingly, although Haspin expression was first identified in testis as a germ cell specific gene (Tanaka et al., 1999), many interrogates about its functions in spermatogenesis are still unsolved. Haspin-GFP in mouse oocytes showed an inconclusive pattern of distribution (Nguyen et al., 2014), but immunolocalization assays detected this kinase through the entire chromatin of condensed bivalents during meiosis-I (Wang et al., 2016). In mouse oogenesis, Haspin is implicated in H3T3 phosphorylation, chromosome condensation, CPC localization at centromeres, microtubule-organizing center (MTOCs) clustering and stability through Aurora C kinase (AURKC) and cytokinesis (Balboula et al., 2016; Nguyen et al., 2018; Quartuccio et al., 2017; Wang et al., 2016). Chemical inhibition of Haspin by 5-Itu causes cell instability and clustering defects of acentriolar MTOCs in meiosis-I (Balboula et al., 2016). This evolutionary conserved functions of Haspin kinases in mammals have recently been also reported in pig oocytes (Cao et al., 2019). However, much fewer reports have been presented for spermatogenesis, where recent data have suggested that the unique spatio-temporal pattern of histone H3 modifications implicates Haspin in the epigenetic control of spermiogenesis (Soupsana et al., 2021).

Since Aurora kinases (AURKs) are the main regulators of chromosome segregation and cytokinesis, it is essential to unravel the signaling pathways that recruit them to the centromeres during gametogenesis. In mammalian germ cells, two different AURKs (AURKB and AURKC) participate in the CPC (Balboula and Schindler, 2014; Shuda et al., 2009), and both can functionally regulate each other’s localization and activity in oogenesis (Nguyen et al., 2018), or compensate for one another ensuring successful mammalian spermatogenesis (Wellard et al., 2020). Therefore, it is crucial to reveal the exact implication of Haspin in the Haspin-H3T3ph-CPC pathway in mammalian male meiosis. In this work, we examined the role of Haspin in male mouse meiosis by using two strategies: a chemical inhibition using LDN-192960 in WT spermatocytes, and the analysis of a Haspin knockout mouse model (*Haspin*^*-/-*^). Both experimental approaches pointed to the implication of Haspin kinase activity in the regulation of chromosome congression, the phosphorylation of AURKB/C at the centromeres and the regulation of centrosome clustering during meiosis-I and meiosis-II. In addition, since Haspin inhibition or ablation does not disturb Shugoshin 2 (SGO2) localization to centromeres, our results suggest that Haspin-H3T3ph-AURKB/C is not the only route to recruit the CPC components to the meiotic centromeres.

## MATERIAL AND METHODS

### Materials

Testes from adult C57BL/6 (wild-type, WT) and genetically modified *Haspin*^*-/-*^ male mice were used for this study. All animal procedures were approved by local and regional ethics committees (Institutional Animal Care and Use Committee and Ethics Committee for Research and Animal Welfare, Instituto de Salud Carlos III) and performed according to the European Union guidelines. The generation of the *Haspin* (*Gsg2*)-null allele, lacking the complete Gsg2 single exon, will be reported elsewhere.

### Culture of seminiferous tubules and LDN-192960 treatment

Culture of seminiferous tubules was performed as previously described (Alfaro et al., 2021; Sato et al., 2011). Testes were removed, detunicated and fragments of seminiferous tubules were cultured in agarose gel half-soaked in MEMα culture medium (Gibco A10490-01) supplemented with KnockOut Serum Replacement (KRS) (Gibco 10828-010) and antibiotics (Penicillin/Streptomycin; Biochrom AG, A2213) with 1 mM LDN-192960 (Sigma, SML0755), and were kept at 34°C in an atmosphere with 5% CO2. Controls were kept in MEMα culture medium without LDN-192916. After 2, 4 and 6 h, control and inhibitor-treated seminiferous tubules were subjected to the squashing technique.

### Immunofluorescence microscopy

Seminiferous tubules were fixed and processed following the technique previously described (Page et al., 1998; Parra et al., 2002). All immunolabellings were performed using a 5% PFA fixation solution.

Squashed seminiferous tubules slides were rinsed three times for 5 minutes in PBS and incubated overnight at 4°C with primary antibodies diluted in PBS. Then, the slides were rinsed three times for 5 minutes in PBS and incubated for 1 hour at room temperature with the secondary antibodies. Finally, the slides were counterstained with 10 µg/ml DAPI for three minutes, rinsed in PBS and mounted with Vectashield (Vector Laboratories).

Kinetochores were revealed with a purified human anti-centromere autoantibody (ACA) serum (Antibodies Incorporated, 435-2RG-7) at a 1:20 dilution. SYCP3 was detected with either a mouse monoclonal antibody against mouse SYCP3 (Santa Cruz Biotechnology, sc-74569) or a rabbit polyclonal antibody recognizing mouse SYCP3 (Santa Cruz Biotechnology, sc-33195) both at a 1:50 dilution. Histone modifications were detected with the following primary antibodies: rabbit polyclonal anti-H2AT120ph antibody (Active Motif, 39391) at a 1:10 dilution, rabbit polyclonal anti-H3T3ph antibodies (Abcam, ab-17532; and Upstate, 07-424) at a 1:800 dilution. AURKB/Cph were revealed with a rabbit polyclonal anti-AuroraTph antibody (Cell Signaling, 2914S) at a 1:30 dilution. SGO2 was revealed with a rabbit polyclonal anti-SGO2 antibody generated by J. L. Barbero (Gómez et al., 2007). Tubulin was detected with a rat anti-α-Tubulin antibody (Abcam, ab-6160) at a 1:100 dilution. Pericentrin (PCNT) was detected with a rabbit polyclonal antibody (Abcam, ab-4448) at a 1:30 dilution. Corresponding secondary antibodies were used against rabbit, mouse, rat and human IgGs conjugated with either Texas Red (Jackson ImmunoResearch), Alexa 594 and Alexa 488 (Molecular Probes). All of them were used at a 1:100 dilution.

After several tries with three different commercial antibodies (Santa Cruz sc-98.622, Abnova H00083903-B01P, Bethyl A302-241A) and two custom-made antibodies, we could not obtain an antibody that accurately show the distribution of Haspin in mouse spermatocytes.

Immunofluorescence images and stacks were collected on an Olympus BX61 microscope equipped with epifluorescence optics, a motorized Z-drive, and an Olympus DP71 or DP70 digital cameras controlled by analySIS software (Soft Imaging System). Finally, images were processed with ImageJ (National Institute of Health, USA; http://rsb.info.nih.gov/ij) or/and Adobe Photoshop softwares.

### Histology

For histological sections, testes were fixed in 10% buffered formalin (SigmaAldrich) and embedded in paraffin wax. After standard washes and dehydratation, Paraplast-embedded tissue blocks were cut in 3-5 μm thick sections in a Reichert microtome. Finally, sections were stained with Hematoxylin/Eosin stain.

### Quantification and statistical analysis

Quantification of the immunofluorescence intensity was estimated by measuring the integrated fluorescence density in individual nuclei using ImageJ by creating a binary mask with the DAPI staining. A minimum of 15-30 metaphase cells per condition were analyzed in each experiment. All graphics and statistical tests were performed with GraphPad Prism 9.0 software. Mean differences for each group were evaluated by an independent sample unpaired *t-*test. Values were expressed as mean ± SEM and p-values below 0.05 were considered statically significant.

## RESULTS

### LDN-192960 is an efficient Haspin inhibitor in mouse organotypic seminiferous tubules cultures

The distribution of the main target of Haspin kinase, H3T3ph, was analyzed by immunolabelling in spermatocytes obtained from cultured seminiferous tubules, using the squashing protocol. This technique does not disturb either the tridimensionality of cell morphology, chromosome condensation or protein distribution in prophase I and dividing spermatocytes (Page et al., 1998; Parra et al., 2002). Labelling of SYCP3 protein, the main component of the synaptonemal complex lateral elements, and DAPI staining were used as markers for the identification of meiosis substages, as previously reported (Parra et al., 2004). H3T3ph begins to be detectable at diakinesis, presenting a heterogeneous faint distribution over the chromatin, slightly more accumulated at chromocenters (clustered centromeres) (Fig. 1 A-B). From prometaphase-I to metaphase-I (M-I from now onwards), H3T3ph intensity progressively rises over the chromatin, being more intense at the interchromatid domain (Fig. 1 C-D). In anaphase-I (A-I), H3T3ph signal becomes weaker (Fig. 1 E) and it completely disappears at the end of telophase-I (T-I) (Fig. 1 F). H3T3ph is not detected in interkinesis (Fig. 1 G) but is again visible at the centromeric heterochromatin at prometaphase-II (Fig. 1 H) and marks completely the chromosomes from metaphase-II (M-II) until the end of telophase-II (Fig. 1 I-K). These data confirm that H3T3ph distribution in mouse male meiosis resembles the pattern described for mitosis, where it was also located over the entire chromatin (Dai and Higgins, 2005).

**Figure 1.**
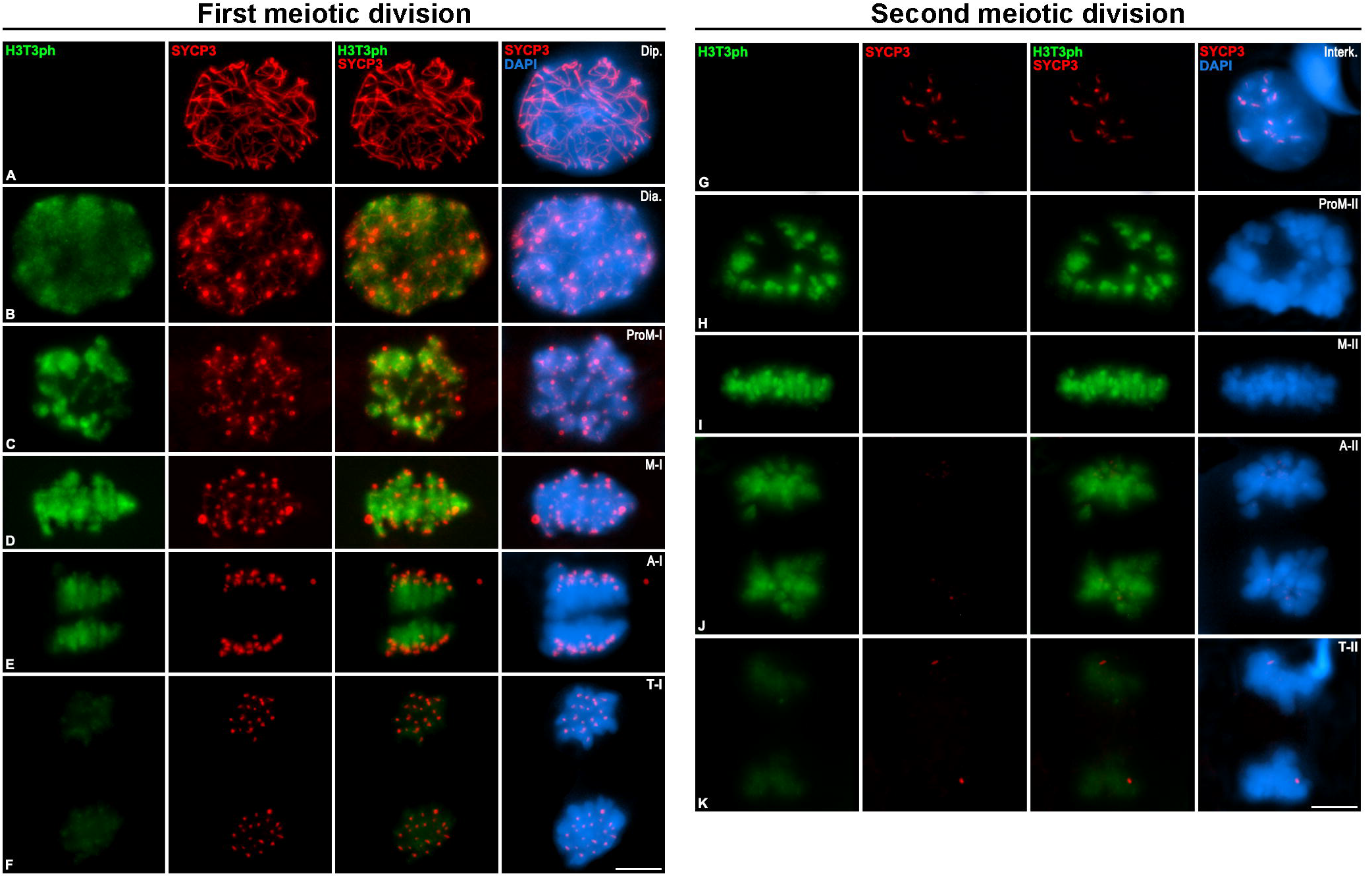
Distribution of H3T3ph and SYCP3 during first and second meiotic division. Double immunolabelling of H3T3ph (green) and SYCP3 (red) on squashed WT mouse spermatocytes on diplotene, (B) diakinesis, (C) prometaphase-I, (D) metaphase-I, (E) anaphase-I, (F) telophase-I, (G) interkinesis, (H) prometaphase-II, (I) metaphase-II, (J) anaphase-II and (K) telophase-II. Chromatin has been stained with DAPI (blue). Scale bar in K represents 10 µm.

We then analyzed the effect of Haspin inhibition *in vitro*. LDN-192960 (LDN) has shown inhibitory effects on the proliferation of somatic cells due to its action over the kinase activity of Haspin (Amoussou et al., 2018; Huertas et al., 2012; Wang et al., 2012). To investigate if this molecule shows activity in spermatocytes, we treated organotypic cultures of seminiferous tubules with increasing concentrations of LDN at different incubation times, using the Haspin target H3T3ph as a marker (Dai and Higgins, 2005; Dai et al., 2005).

In somatic cell lines (HeLa and U2OS) the optimal efficient concentration used was 10 μM (Wang et al., 2012). However, we had to use higher concentrations. This is due to the peculiarities of the organotypic culture, in which tubules are placed over an agarose gel (half-soaked) embedded in the medium. This means that the media does not surround the cells, but instead, the media diffuses to the seminiferous tubules via the agarose gel (Alfaro et al., 2021; Sato et al., 2011). We therefore decided to test higher concentrations of the inhibitor for 2, 4 and 6 hours. Concentrations below 1 mM had no effect on meiosis progression or cell morphology, but conspicuous effects were observed with a 1 mM concentration (Figure 2). These effects involved both alterations of the morphology of cells and the intensity of H3T3ph signal in M-I spermatocytes. Results showed that 13.1% of M-I cells lost H3T3ph signal after 2 hours of treatment (Fig 2a A-B, 2b and 2c). In contrast, after 4 hours of LDN treatment, we observed that H3T3ph signal was abolished in 97.9% of M-I cells (Fig 2a C-D and 2b,c). The total loss of H3T3 phosphorylation in M-I cells was achieved after 6 hours LDN treatment (Fig 2a E-F and 2b,c). These low measurements of intensity correspond to the background signal. These results confirmed that the Haspin kinase activity over H3T3 is abolished in mouse spermatocytes with 1 mM LDN 6h treatment. For this reason, all the subsequent analyses were performed using these culture conditions.

**Figure 2.**
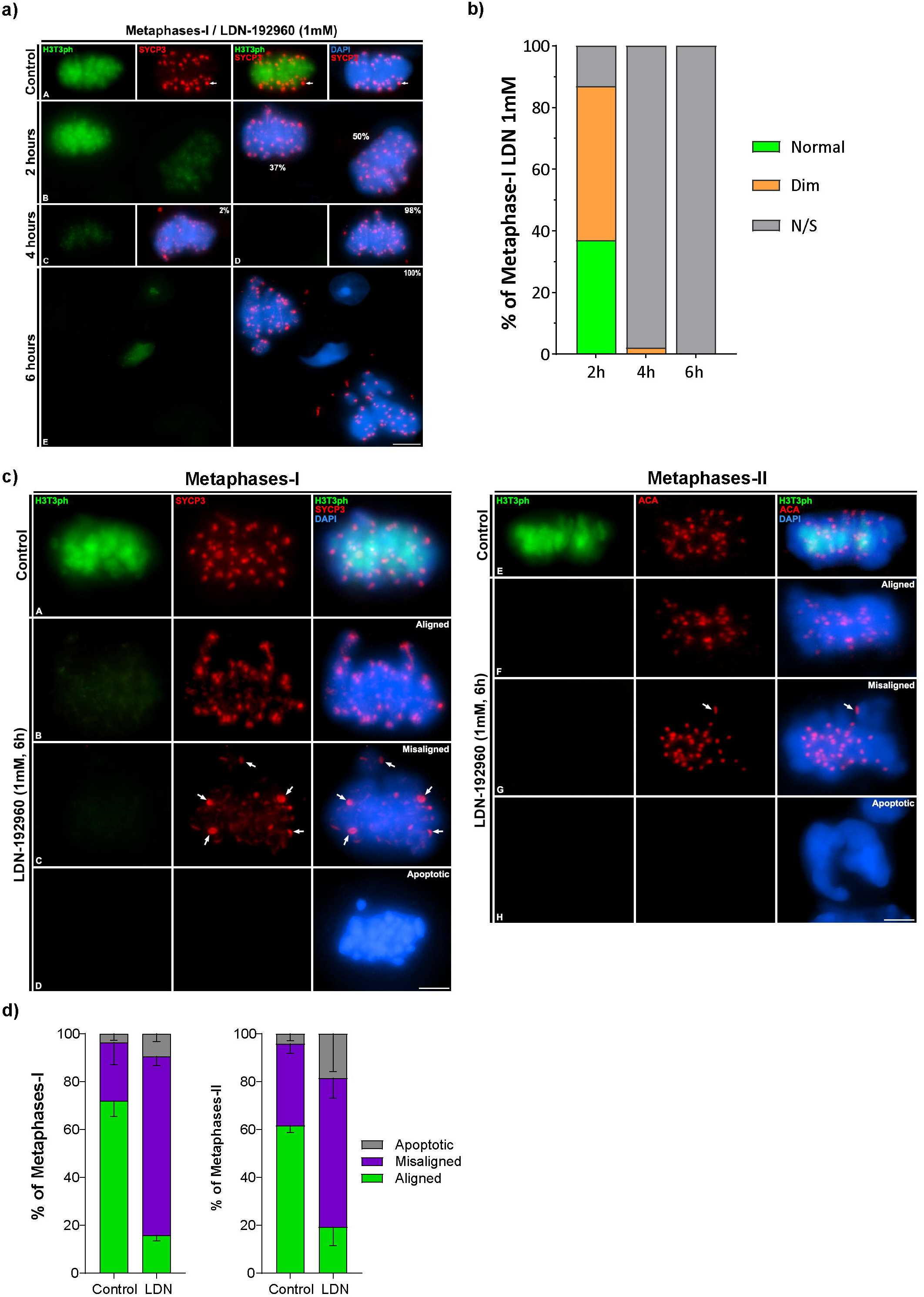
Optimisation of Haspin inhibition in organotypic cultures of seminiferous tubules. a. Double immunolabelling of H3T3ph and SYCP3 in squashed spermatocytes for (A) control and (B) 2h, (C, D) 4h and (E) 8h 1 mM of LDN-192960. Scale bar in E represents 10 µm. b. Quantification of metaphase-I percentage with H3T3ph signal at different timings in cultured spermatocytes with 1 mM of LDN-192960. Data represents the percentage of metaphase-I with non-signal, dim or normal signal of H3T3ph for 2h, 4h and 8h of treatment. c. H3T3ph distribution in control and 1mM 6h LDN-192960-treated spermatocytes. Double immunolabelling of H3T3ph (green) and SYCP3 (red) in (A) control and (B-D) 1mM 6h LDN-192960-treated metaphase-I. Double immunolabelling of H3T3ph (green) and ACA (red) in (E) control and (F-H) 1mM 6h LDN-192960-treated metaphase-II. Chromatin has been stained with DAPI (blue). Scale bar in D and H represents 10 µm. d. Quantification of the incidence of misaligned metaphase-I and metaphase-II. Percentage of aligned, misaligned and apoptotic metaphases is represented for control and 1 mM LDN-192960 for 6h treatment. Experiments were conducted in three different biological replicates. Bars and error bars represent mean ± SD.

### Haspin activity is required for proper chromosome congression in M-I and M-II mouse spermatocytes

The effect of LDN were evident not only over the phosphorylation of H3T3, but also on the alignment of chromosomes at the metaphase plate. In order to better characterize this effect, we performed a quantitative analysis of the phenotypes showed by M-I and M-II spermatocytes after labeling H3T3ph and SYCP3 (for M-I) and H3T3ph and centromeres (for metaphase-II, M-II). Cells were classified in three categories: aligned M-I or M-II (all bivalents/chromosomes are arranged at the equatorial plate), misaligned M-I or M-II (one or more bivalents/chromosomes are not in the equatorial plate) and apoptotic M-I or M-II (chromatin hypercondensation and no immunoreactivity) (Fig. 2c). H3T3ph signal is clearly detected in control M-I (Fig. 2c A) and M-II (Fig 2c E). In contrast, H3T3ph signal is absent in all LDN-treated M-I and M-II cells, regardless they were classified as aligned or misaligned (Fig. 2c). Apoptotic cells were not immunoreactive, although their size and morphology allowed them to be identified as M-I (Fig. 2c D) or M-II (Fig. 2c H).

The quantitative analysis showed that LDN-treated cultures showed a 3-fold increase in chromosome misalignment in M-I spermatocytes (74.7%) compared with control cells (24.3%) and a 2-fold increase in apoptotic rate (9%) (Fig. 2d). Similarly, the percentage of misaligned M-II rise 2-fold in LDN-treated spermatocytes (62.3%), compared with control (34.2%). For M-II, the apoptotic rate in LDN-treated spermatocytes (18.4%) was 4-fold higher than in control ones (4.1%) (Fig. 2d). These data indicate that the disruption of Haspin kinase activity causes a dramatic increase in alignment errors during M-I and M-II in mouse spermatocytes, potentially leading to cell death.

### Haspin inhibition perturbs kinetochore-microtubule attachment and centrosome clustering in mouse spermatocytes

Since LDN treatment affects M-I and M-II chromosome congression, we then studied the interaction between kinetochores and the meiotic spindle. Haspin involvement in MTOCs organization has been previously reported in oocytes (Balboula et al., 2016). We therefore studied the dynamics of the meiotic spindle and the attachment of chromosomes to microtubules. For this purpose, we immunolabelled spermatocytes with α-Tubulin and a ACA serum that labels the kinetochores (Fig. 3). M-I with correctly bioriented and aligned bivalents had bipolar spindles (Fig. 3a A). In contrast, in misaligned M-I some bivalents were not correctly anchored to the microtubules, usually remaining linked to only one pole (monothelic orientation) (Fig. 3a B). A careful examination of these misaligned bivalents revealed that the two homologous kinetochores were usually attached to the same cellular pole (Fig 3b B′ and B″). No apparent errors in cohesion were observed as univalents or independent chromatids were never observed. Surprisingly, M-I with two or more misaligned bivalents usually presented tripolar spindles (Fig. 3a C). The incidence of multipolar spindles in LDN-treated spermatocytes raised to 10,88% in M-I and 6,25% in M-II (Fig. 3c). These data suggest a possible impairment in the centrosome dynamics. Therefore, we then performed an immunolocalization of Pericentrin (PCNT), the integral component of the pericentriolar matrix of mouse centrosomes. In control M-I, PCNT appears as a compact mass in both poles of the cell (Alfaro et al, 2021). However, LDN-treated cells presented alterations in PCNT distribution. In bipolar spindles, the signal appeared disaggregated showing a variable number of pericentriolar matrix clusters with an irregular morphology (Fig. 3d A-B). These features were also obvious in multipolar M-I or M-II cells (Fig. 3d C-E).

**Figure 3.**
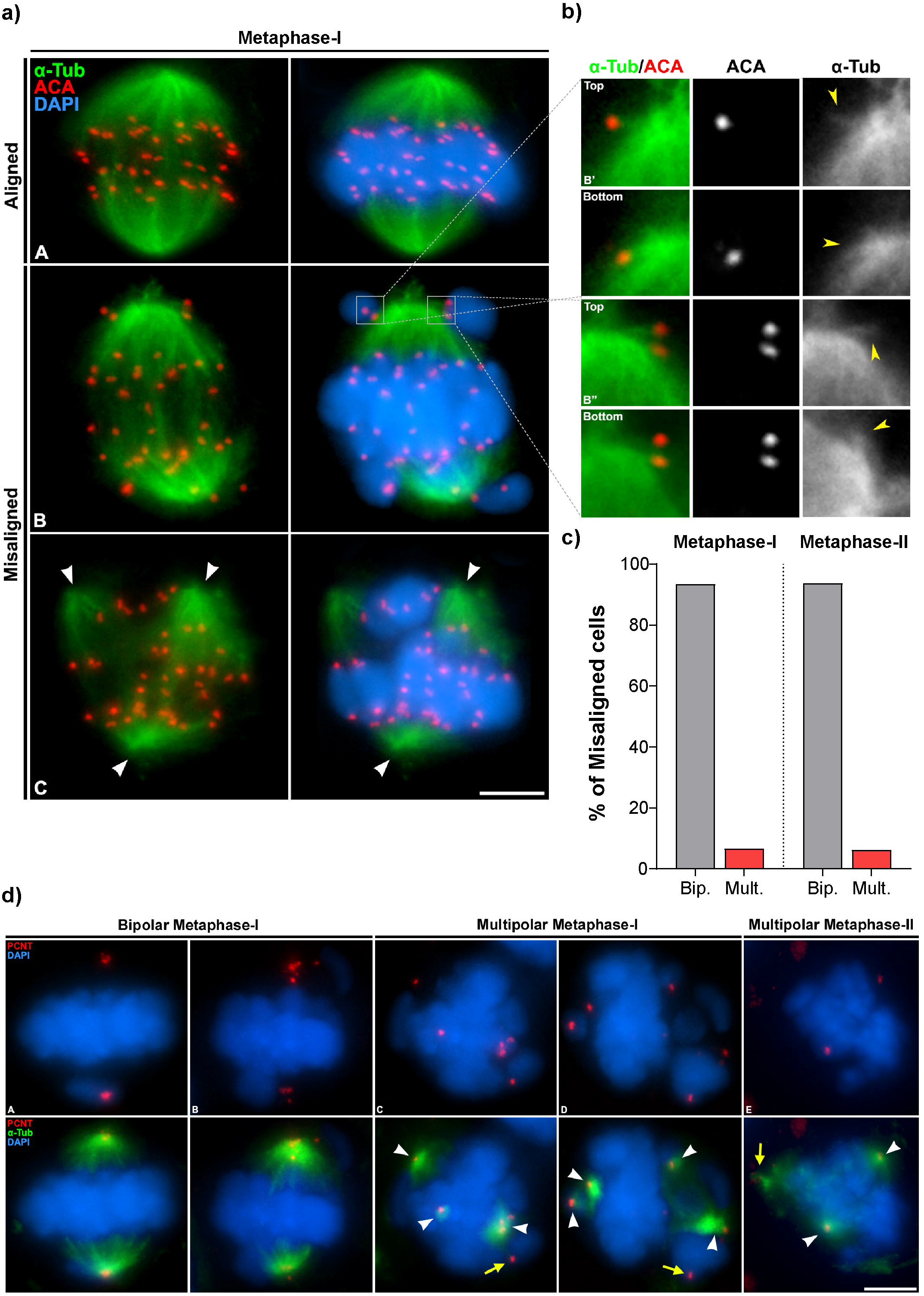
Analysis of chromosome congression in metaphase-I and metaphase-II after in 1mM 6h LDN-192960 treatment. a. Double immunolabelling of α-Tubulin (green) and ACA (red) in aligned (A) and misaligned (B, C) metaphase-I. Chromatin has been stained with DAPI (blue). White arrows in C indicate the location of spindles. Scale bar in C represents 10 µm. b. Centromere attachment of misaligned bivalents. Double immunolabelling of α-Tubulin (green) and ACA (red). Each inset corresponds to the centromere of each homologous chromosomes (top and bottom). Yellow arrows correspond with microtubule-centromere attachment. Chromatin has been stained with DAPI (blue). Scale bar in C represents 10 µm. c. Quantification of the percentage of multipolar spindles during first and second meiotic division. d. Double immunolabelling of α-Tubulin (green) and PCNT (red) of bipolar spindle metaphase-I and multipolar spindle metaphase-I and metaphase-II. Chromatin has been stained with DAPI (blue). Scale bar in E represents 10 µm.

### AURKB/C phosphorylation is reduced at centromeres when H3T3 phoshorylation is abolished

The phosphorylation of the H3T3 has been considered critical for the recruitment of the CPC (Higgins, 2010; Wang et al., 2010). Therefore, the downstream AURKB kinase recruitment is directly linked to Haspin kinase activity (Krenn and Musacchio, 2015; van der Horst et al., 2015). AURKA, AURKB and AURKC belong to a family of kinases with high sequence homology (Tang et al., 2017). AURKA participates in centrosomes dynamics in male mouse meiosis (Alfaro et al., 2021; Wellard et al., 2021), while AURKB and AURKC (hereafter denoted collectively as AURKB/C) function as the catalytic subunit of the CPC, presenting overlapping functions in spermatogenesis (Jordan et al 2021).

We used an antibody that detects these three kinases in their phosphorylated active form: phosphorylated Aurora kinases (AURKph). In control spermatocytes, this antibody mainly labelled centromeres during M-I and M-II, as well as centrosomes and the midbody at TI and TII (Figure S1). According to previous reports, these marks correspond to the localization of phosphorylated AURKB/C (AURKB/Cph) at centromeres (Nguyen and Schindler, 2017) and phosphorylated AURKA at the centrosomes (AURKAph) (Alfaro et al., 2021; Tang et al., 2017), respectively. As shown previously, H3T3ph decorated the chromatin in control MI (Fig. 4a A) and MII (Fig. 4b A), but this histone modification was absent when Haspin kinase was inhibited with LDN in both MI (Fig. 4a B) and MII (Fig. 4b B). After Haspin inhibition, the recruitment of AURKB/Cph to the ICD in M-I (Fig. 4a D) and in M-II (Fig. 4b D) is altered, as the intensity of labeling is fainter than in control spermatocytes (Fig. 4a C and 4b C). Quantification of the fluorescence signals showed a significant decrease in AURKB/Cph intensity at the centromeres both in M-I (Fig. 4c) and M-II (Fig. 4d). Altogether, these data suggest the implication of Haspin kinase activity in the AURKB/C phosphorylation at the centromeres. Surprisingly, the signals at the spindle poles in LDN-treated spermatocytes were similar to that found in control spermatocytes (white arrowheads in Fig. 4a and 4b), suggesting that AURKAph might not be disturbed.

**Figure 4.**
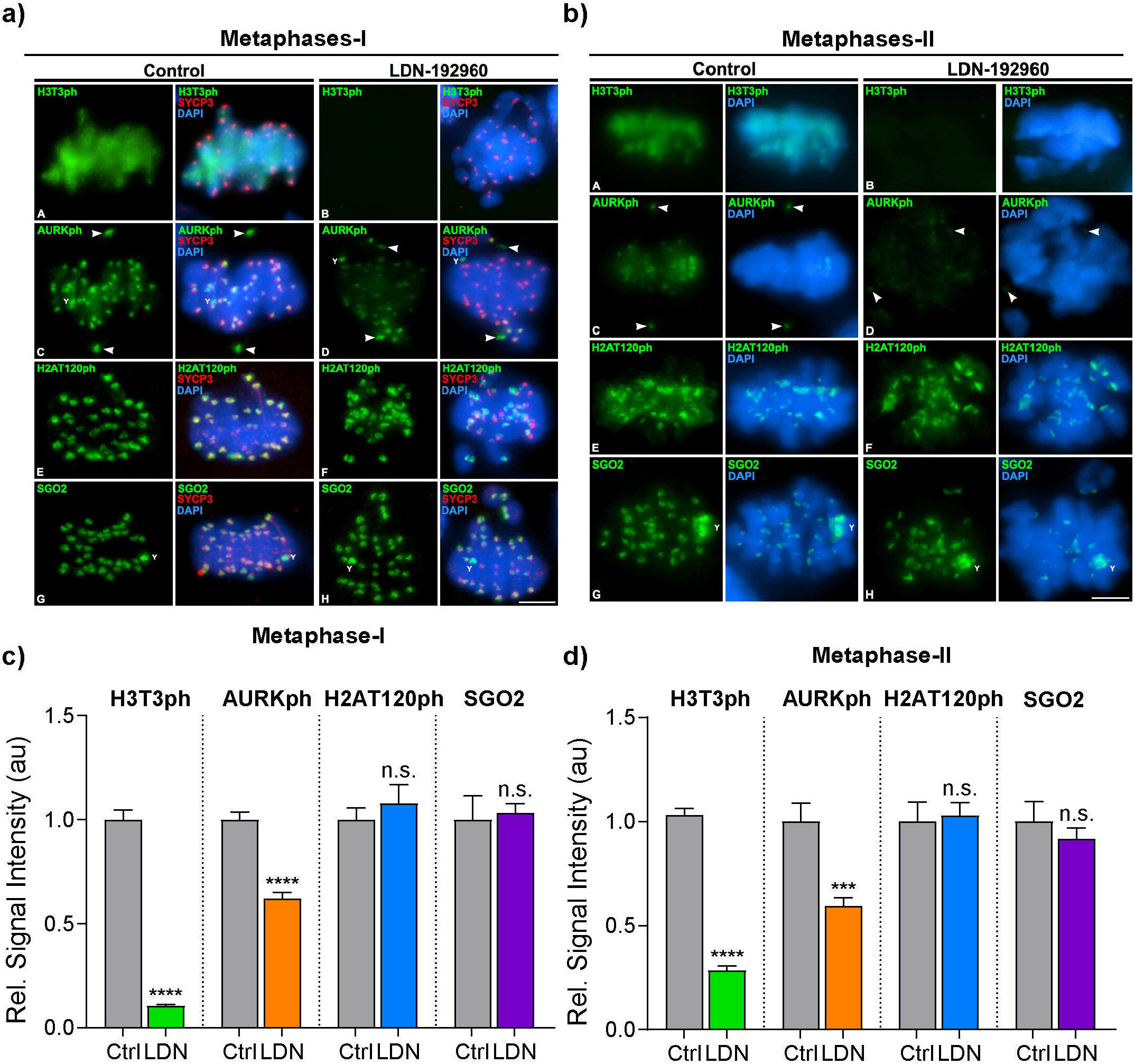
Analysis of the Inner Centromere Domain (ICD) in 1mM 6h LDN-192960-treated spermatocytes. a. ICD components distribution during metaphase-I spermatocytes. Double immunolabelling of (A-B) H3T3ph (green) and SYCP3 (red), (C-D) AURKph (green) and SYCP3 (red), (E-F) H2AT120ph (green) and SYCP3 (red), and (G-H) SGO2 (green) and SYCP3 (red). Chromatin has been stained with DAPI (blue). Scale bar in H represents 10 µm. b. ICD components distribution during metaphase-II spermatocytes. Double immunolabelling of (A-B) H3T3ph (green) and SYCP3 (red), (C-D) AURKph (green) and SYCP3 (red), (E-F) H2AT120ph (green) and SYCP3 (red), and (G-H) SGO2 (green) and SYCP3 (red). Chromatin has been stained with DAPI (blue). Scale bar in H represents 10 µm. c. Quantitative analysis of ICD components signals in metaphase-I spermatocytes. Experiments were conducted for one biological replicates. Data represents mean ± SEM, **** *p* < 0.0001, Student’s *t*-test. d. Quantitative analysis of ICD components signals in metaphase-II spermatocytes. Experiments were conducted for one biological replicates. Data represents mean ± SEM, **** *p* < 0.0001, Student’s *t*-test.

### H3T3ph absence does not alter SGO2 loading to centromeres

Given the likely implication of Haspin in AURKB/C phosphorylation at centromeres, we then wondered if SGO2, the protector of meiotic centromeric cohesion, could be affected. In mitosis, recruitment of the Shugoshins to centromeres requires the phosphorylation of H2AT120 by the kinetochore kinase Bub1 (Kawashima et al., 2010). With these premises in mind, we then studied the distribution of the meiotic centromere cohesion regulator SGO2 and its centromere docking mark H2AT120ph.

H2AT120ph appears localized to the ICD of all chromosomes in both control (Fig. 4a E) and LDN-treated M-I (Fig. 4a F). Similarly, SGO2 is present in the ICD of all chromosomes in control M-I (Fig. 4a G) and also LDN-treated M-I (Fig. 4a H). Similarly, signals of H2AT120ph and SGO2 in control and LDN-treated M-II are consistent in distribution and intensity (Fig. 4b E-H). H2AT120ph appeared more concentrated at the centromere of the Y chromosome both in control and treated MI and MII (Fig. 4a G-H and Fig. 4b G-H). Quantification analysis showed no significant differences in H2AT120ph nor SGO2 intensity at the centromeres between control and LDN inhibition in M-I (Fig. 5c) and M-II (Fig. 4d). These data suggest that the kinase activity of Haspin does not have a direct role in the maintenance of the H2AT120ph-SGO2 pathway nor in the centromeric cohesion in male mouse meiosis.

**Figure 5.**
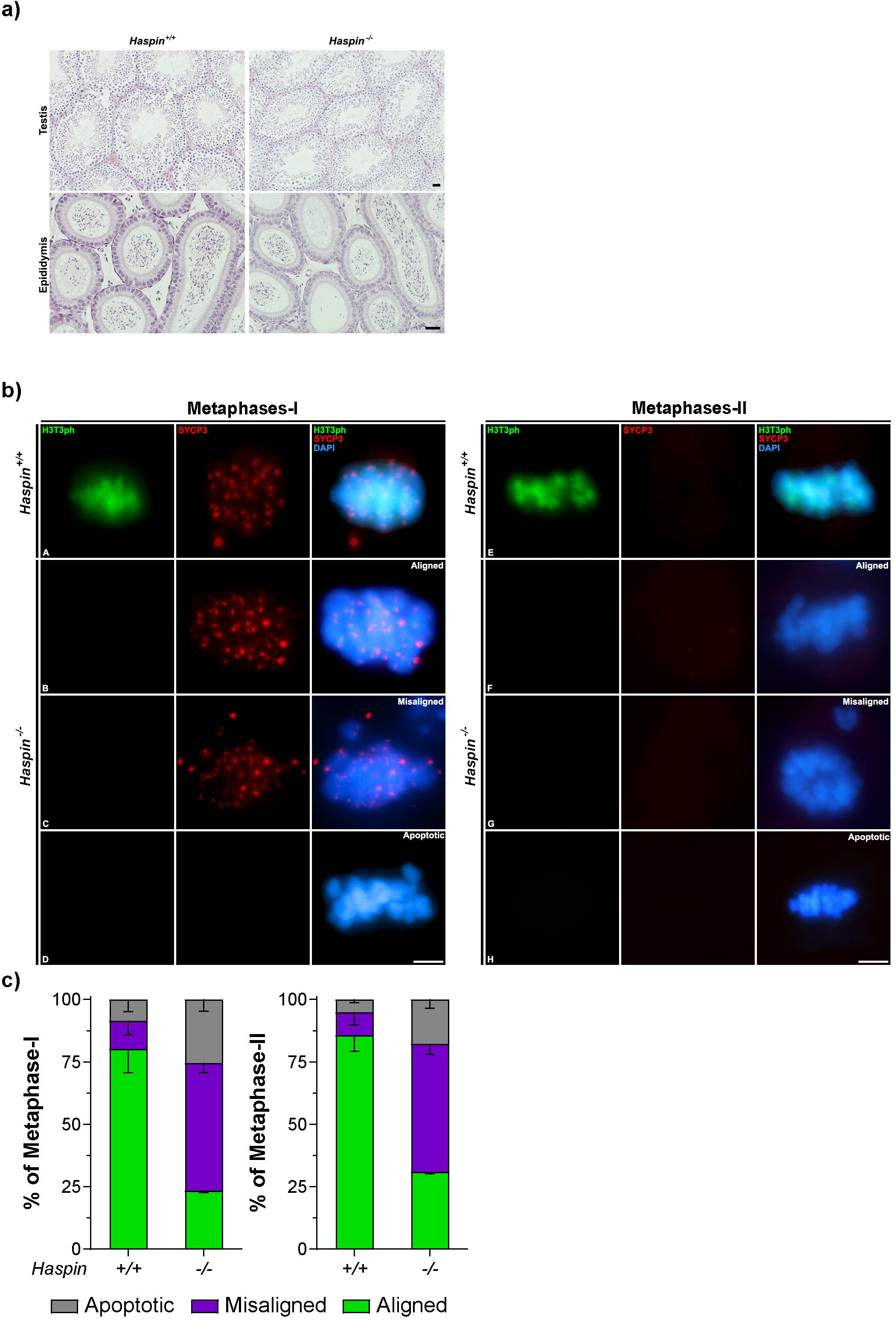
Analysis of the meiotic phenotype of *Haspin* knockout mouse model. a. Histological sections of seminiferous tubules and epididymis of *Haspin*^*+/+*^ *and Haspin*^*-/-*^. Scale bar represents 200 µm. b. Distribution of H3T3ph in *Haspin*^*+/+*^ *and Haspin*^*-/-*^ spermatocytes. Double immunolabelling of H3T3ph (green) and SYCP3 (red) in (A) *Haspin*^*+/+*^ and (B-D) *Haspin*^*-/-*^ metaphase-I. Double immunolabelling of H3T3ph (green) and SYCP3 (red) in (E) *Haspin*^*+/+*^ and (F-H) *Haspin*^*-/-*^ metaphase-II. Chromatin has been stained with DAPI (blue). Scale bar in D and H represents 10 µm. c. Quantification of the incidence of misaligned metaphase-I and metaphase-II. Percentage of aligned, misaligned and apoptotic metaphase is represented for *Haspin*^*+/+*^ *and Haspin*^*-/-*^. Experiments were conducted in two different biological replicates. Bars and error bars represent mean ± SD.

### *Haspin*^*-/-*^ individuals are fertile but spermatocytes lack centromeric H3T3ph and show alterations in chromosome alignment

To corroborate the *in vitro* results, we then analyzed the meiotic phenotype of a *Haspin* knockout (KO) mouse model (*Haspin*^*-/-*^) (Fig. 5). Haspin-KO mitotic cell lines have been described to present delays in metaphase-anaphase progression in mitosis due to bi-orientation errors and a weakness of centromeric cohesion (Zhou et al., 2014). However, *Haspin*^*-/-*^ fibroblasts did not show either severe mitosis progression delay or apparent centromeric cohesion alterations (data not shown).

We analyzed the histology of the testis of the mouse model *Haspin*^*-/-*^. Seminiferous tubules and the peritubular tissue of *Haspin*^*-/-*^ seemed completely normal and spermatozoa could be observed in the epididymis (Fig. 5a). This suggests an apparently normal fertility of these mutants, which was confirmed after crossing of Haspin-deficient males or females with wild-type mice. Both female and male *Haspin*^*-/-*^ individuals are fertile producing normal litter sizes.

In agreement with the *in vitro* experiments with LDN inhibitor, we observed that H3T3ph is absent in the chromosomes of both M-I and M-II spermatocytes in *Haspin*^*-/-*^ mice (Figure 5b). Likewise, we reported an abundance of misaligned M-I and M-II. The frequency of these misaligned metaphases was even higher than that reported in our *in vitro* experiments (51,13% in M-I and 51.21% in M-II) (Fig 5d). Accordingly, the frequency of apoptotic metaphases increased 3-fold during M-I (25.42%) compared to WT (8.5%), while LDN-treatment was only 2-fold higher than in control. However, the ratio of *Haspin*^*-/-*^ apoptotic M-II (18%) was similar, but almost 4-fold higher than in WT (5.1%) (Fig 5c). Surprisingly, multipolar metaphases were not observed.

We finally analyzed the location of the ICD components in *Haspin*^*-/-*^ spermatocytes. We observed an almost complete absence of H3T3ph in *Haspin*^*-/-*^ M-I (Fig. 6a A-B and 6c)) and M-II (Fig. 6b A-B and 6c)). The absence of H3T3ph is accompanied by a significant reduction of AURKB/Cph at centromeres in MI (Fig. 6a C-D and 6c) and MII (Fig. 6b C-D and 6d). In contrast, both H2AT120ph and SGO2 abundance was unaltered in MI (Fig. 6a E-F and 6c) and MII (Fig. 6a E-F and 6d) Quantification analysis showed no significant differences in H2AT120ph nor SGO2 intensity at the centromeres between *Haspin*^*+/+*^ and *Haspin*^*-/-*^ in M-I (Fig. 6c) and M-II (Fig. 6d).

**Figure 6.**
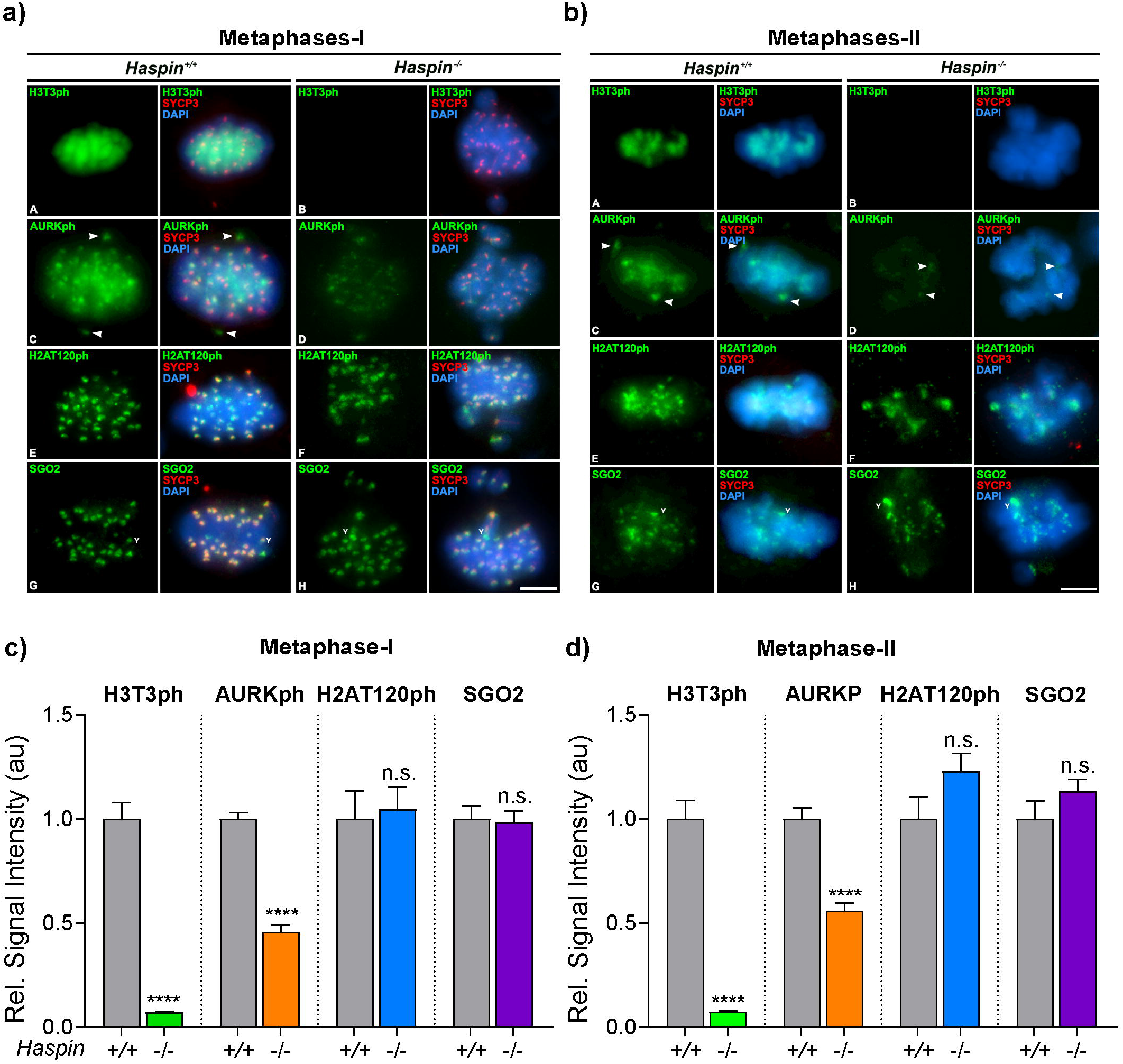
Analysis of the Inner Centromere Domain (ICD) in *Haspin*^*+/+*^ *and Haspin*^*-/-*^spermatocytes. a. ICD components distribution during metaphase-I. Double immunolabelling of (A-B) H3T3ph (green) and SYCP3 (red), (C-D) AURKph (green) and SYCP3 (red), (E-F) H2AT120ph (green) and SYCP3 (red), and (G-H) SGO2 (green) and SYCP3 (red). Chromatin has been stained with DAPI (blue). Scale bar in H represents 10 µm. b. ICD components distribution during metaphase-II spermatocytes. Double immunolabelling of (A-B) H3T3ph (green) and SYCP3 (red), (C-D) AURKph (green) and SYCP3 (red), (E-F) H2AT120ph (green) and SYCP3 (red), and (G-H) SGO2 (green) and SYCP3 (red). Chromatin has been stained with DAPI (blue). Scale bar in H represents 10 µm. c. Quantitative analysis of ICD components signals in metaphase-I spermatocytes. Experiments were conducted for two biological replicates. Data represents mean ± SEM, **** *p* < 0.0001, Student’s *t*-test. d. Quantitative analysis of ICD components signals in metaphase-II spermatocytes. Experiments were conducted for two biological replicates. Data represents mean ± SEM, **** *p* < 0.0001, Student’s *t*-test.

## DISCUSSION

The role of Haspin has been widely investigated in somatic cells. This protein was found to be responsible of H3T3 phosphorylation, which is a necessary step for the proper loading of the CPC to the centromeres, chromosome alignment and sister chromatid cohesion (Dai and Higgins, 2005; Maiolica et al., 2014; Markaki et al., 2009). Here, we report that some of these functions are also conserved in meiosis, although important differences are found, indicating a possible redundancy of pathways in the correct assembly of centromeres in meiosis. The two approaches used here, the functional *in vitro* inhibition with LDN and the use of a knockout mouse model produced congruent results, indicating the accuracy of these methods to assess Haspin function in male meiosis.

### Haspin ensures proper chromosome congression in male mouse meiotic divisions by phosphorylating H3T3 at centromeres, but is dispensable for the loading of SGO2 to the ICD

In the past years, several reports suggested that the CPC, including AURKB, must locate to the ICD to ensure chromosome congression (Hindriksen et al., 2017; Kelly et al., 2010; Santaguida et al., 2010; Saurin et al., 2011; Wang et al., 2010). The current model proposes that the CPC is recruited to the chromatin region where H3T3ph and H2AT120ph are present (Yamagishi et al., 2010), implying that the ICD may be defined by simultaneous interactions of the CPC with H3T3ph, H2AT120ph and SGO1/2 (Hadders et al., 2020; Krenn and Musacchio, 2015; Liang et al., 2020). Interestingly, a recent model suggested that the ICD is scaffolded by an spatially regulated CPC, that it able to assembly and disassembly allowing histone modifications to interact with each other during the different stages of the cell cycle (Trivedi and Stukenberg, 2020). In this regard, several publications support the idea of different routes to recruit AURKB to mitotic centromeres. Broad and coworkers investigated three discrete AURKB populations at mitotic centromeres: an inner centromere pool of AURKB would be recruited by Haspin phosphorylation of histone H3; a second pool, located at the kinetochores, would be recruited by Bub1 phosphorylation of histone H2A; a third pool would be recruited to the outher kinetochore by protein CENP-C in early mitosis independently of either Bub1-H2AT120ph-SGO1 or Haspin-H3T3ph pathways (Broad et al., 2020). Furthermore, two additional publications demonstrated that either Haspin or Bub1 kinase activity are sufficient to recruit AURKB to mitotic centromeres (Hadders et al., 2020; Liang et al., 2020), hence suggesting that Bub1-H2AT120ph and Haspin-H3T3ph coexist in the ICD and have combined actions that ensure different pools of AURKB to the centromeres. Finally, another recent report argued that in oocytes Helicase LSH could also be responsible for H3T3 phosphorylation, suggesting a close relationship between chromatin ultrastructure, transcriptional activity and histone modifications at centromeres (Baumann et al., 2020).

The results presented here are congruent with the involvement of Haspin in the proper assembly of centromeres during meiosis and interestingly also suggest a redundancy of pathways in some steps of this process. Centromere dynamics is clearly altered in the absence (*Haspin*^*-/-*^ spermatocytes) or inhibition (LDN-treated spermatocytes) of Haspin kinase activity. The most conspicuous effects are the abolition of H3T3ph signal and the reduction of AURKph signal at centromeres of both meiotic divisions. These findings suggest a clear implication of Haspin kinase activity in the AURKB/C phosphorylation pathway at the centromeres. This probably leads to a malfunction of centromeres, as revealed by the increase of misaligned chromosomes and apoptotic cells in both M-I and M-II. Therefore, Haspin may play a fundamental role in ensuring proper chromosome congression/segregation.

However, this role may be redundant with other proteins. We found that neither Haspin absence (*Haspin*^*-/-*^) nor inhibition (LDN treatment) alter SGO2 localization to meiotic centromeres. This points to an alternative route capable of recruiting SGO2 to the ICD in order to fulfill its function as centromeric cohesion protector (Gómez et al., 2007; Llano et al., 2008). In mitosis, AURKB phosphorylates SGO2 (Tanno et al., 2010). Moreover, the inhibition of Bub1 kinase activity in Haspin-deficient cells abolished any detectable enrichment of AURKB at centromeres (Liang et al., 2020). The few data available about mammalian gametogenesis indicate that in mouse oocytes the kinase activity of Bub1 is dispensable for SGO2 localization at centromeres (El Yakoubi et al., 2017). Here, we show that albeit Haspin kinase is responsible for the recruitment of phosphorylated AURKB/C at centromeres, abolition of the Haspin-H3T3ph pathway does not preclude the recruitment of SGO2 to the ICD. Consequently, we propose that in mouse spermatogenesis Bub1-H2AT120ph-SGO2 and Haspin-H3T3ph-CPC pathways might also have redundant functions in the assembly of the CPC at centromeres. The fact that *Haspin*^*-/-*^ mice are fertile despite of the absence of H3T3ph and the reduction of AURKB phosphorylation at centromere support this hypothesis. Moreover, given that either the Haspin or the Bub1 kinase could independently recruit different pools of Aurora to support faithful chromosome segregation in mitosis (Hadders et al., 2020; Liang et al., 2020), we could hypothesize that in spermatogenesis there might also been different pools of AURKB at the inner centromere that synergistically ensure chromosome congression in meiosis. Intriguingly, meiosis possesses a third AURK (AURKC) in comparison to mitosis (Brown et al., 2004; Nguyen and Schindler, 2017). Therefore, given that Haspin inhibition does not perturb the spindle assembly checkpoint activity in Aurkc^−/−^ during female mammalian meiosis I (Quartuccio et al., 2017), and that AURKB and AURKC have collaborative functions in oogenesis (Nguyen et al., 2018) and spermatogenesis (Wellard et al., 2020), we could argue that Haspin-AURKB interaction might be dispensable and compensated by AURKC functions during spermatogenesis. Utterly, all these data point to a Haspin-dependent centromeric AURKB function in the context of chromosome congression regulation that might have synergic functions with centromeric AURKC in male mouse meiosis. Altogether, our work suggests that during spermatogenesis, Haspin-H3T3ph is not the only route to recruit CPC components to the meiotic centromeres and is dispensable for the loading of SGO2 to centromeres in male mouse meiosis. Instead of converging to accumulate the CPC precisely at the inner centromere, Haspin and Bub1 might each also recruit a separate functional CPC pool to the centromere, potentially sharing similarities with the mitotic centromeric pathway (Broad et al., 2020). In conclusion, without excluding that Haspin might have different functions adapted to the specific cell characteristics among mitosis, and between female and male meiosis, we can here deduce that this kinase is dispensable for meiosis.

### Haspin is implicated in centrosome dynamics in male mouse meiosis

Several data suggested over the last years that Haspin inhibition causes MTOC instability in somatic cells and oocytes. In mitosis (U2OS cells), eGFP-Haspin is detected at centrosomes (Dai et al., 2005) and further analysis described that Haspin knockdown with siRNA increased the number of centrosome-like foci that contain γ-Tubulin and allow microtubule polymerization (Dai et al., 2009). On the other hand, an increased incidence of MTOC foci in M-I spindles was also observed while analyzing Haspin functions in mouse oocytes using inhibitors and overexpression approaches (Nguyen et al., 2014). More recent results indicated that after Haspin *in vitro* inhibition, the multiple MTOC foci that should cluster into two poles fail to do so, albeit the MTOC-localization of AURKA was not perturbed (Balboula et al., 2016).

Our *in vitro* results suggest that the inhibition of Haspin with LDN also causes centrosome instability and the appearance of multipolar M-I and M-II spermatocytes, pointing to a potential role of Haspin in centrosome regulation in male mouse meiosis. In these multipolar metaphases all induced MTOCs are capable of recruiting Pericentrin and polymerize microtubules, sharing similarities with somatic cells (Dai et al., 2009) and mouse oocytes (Balboula et al., 2016). These results suggest that Haspin kinase might participate in centrosome regulation, which is consistent with the localization of GFP-Haspin in oocytes at MTOCs (Balboula et al., 2016). Moreover, this hypothesis is reinforced by the identification of some proteins known to localize at mitotic centrosomes that are phosphorylated in a Haspin-dependent manner (Maiolica et al., 2014). Remarkably, we detect AURKAph at the poles of misaligned M-I and M-II in our *in vitro* studies with LDN and in the mouse model *Haspin*^*-/-*^. This suggests that Haspin is not directly implicated in AURKA recruitment to centrosomes and it is not responsible for AURKA phosphorylation at centrosomes in spermatocytes, accordingly to previous data in oocytes (Balboula et al., 2016). We argue that these multipolar metaphases are only detected in the *in vitro* experiments because the inhibition of the kinase activity is abolished abruptly. In contrast, in the mouse model alternative routes could have been stablished to compensate Haspin roles at the centrosome. Nevertheless, we cannot exclude that unknown off targets of LDN could also be affected in the experimental *in vitro* experiments.

### Haspin inhibition or ablation induces chromosomal mis-segregation in male mouse meiosis but does not alter fertility

When Haspin inhibition induces chromosomes mis-segregation in somatic cells, error in congression led to cytokinesis failure, mitotic exit and micronuclei formation (Dai et al., 2009). In contrast, in mouse oocytes, anaphase I proceeds with chromosomes being mis-segregated and persistence of improper kinetochore-microtubule attachments, aberrant chromatin condensation and cohesion abnormalities, yet cells complete cytokinesis and progress to metaphase II carrying aneuploidies (Kang et al., 2015; Nguyen et al., 2014; Quartuccio et al., 2017; Wang et al., 2016). We here found that Haspin LDN-inhibition in spermatocytes leads to a high incidence of misaligned metaphases, suggesting a direct role of this kinase on chromosome congression in both meiotic divisions that could potentially cause aneuploid gametes. The *Haspin*^*-/-*^ *m*ouse model unravels the complexities of fully depleting Haspin kinase *in vitro* experimentally and confirmed Haspin implication in chromosome congression. Nevertheless, *Haspin*^*-/-*^ male mice present sperm in the epididymis, indicating that although a high incidence of mis-segregations errors occur during the first and the second meiotic division, some spermatocytes progress to form mature gametes. Altogether, the combined data regarding Haspin inhibition in mitosis and meiosis is intriguing. The literature discussed shows that Haspin inhibition leads to cell death in mitosis, but it does not interrupt meiosis, yet it needs to be considered that all these data have been conducted *in vitro*, where Haspin is abruptly disrupted and might present a more aggressive phenotype. Our work combines the *in vitro* experiments with the characterization of the meiotic phenotype of a *Haspin*^*-/-*^ mouse model, allowing us to conclude that on the one hand, Haspin is not needed for faithful mitosis, as *Haspin*^*-/-*^ animal health is not disrupted. On the other hand, given than both females and males *Haspin*^*-/-*^ are fertile, we can also conclude that Haspin kinase is dispensable for male mouse fertility.

## ACKOWLEDGEMENTS

We express our sincere thanks to José Luis Barbero for providing antibodies; to Katja Wassmann for scientific support; and to Lorena Barreras for technical assistance. This work was funded by Ministerio de Economía y Competitividad and Ministerio de Ciencia e Innovación (Spain), grant numbers: PID2020-117491GB-I00 to J.A.S.; RTI2018-095582-B-I00 to M.M.; and CGL2014-53106-P to J.P. I.B was supported by grant FPI of Ministerio de Economía, Industria y Competitividad (Spain). C.M. was supported by Juan de la Cierva programme of Ministerio de Economía, Industria y Competitividad (Spain) and by CNIO Friends Postdoctoral Programme (CNIO, Spain).

## AUTHOR CONTRIBUTIONS

IB, JAS and RG conceived the study; IB, PL, IM, CM, AV, MM, JAS and RG performed all experiments; IB, PL, IM, JP, JAS and RG analyzed results; IB, PL and RG wrote the paper with some contributions from the other co-authors.

## Legend of Figures

**Supplementary Figure 1.**
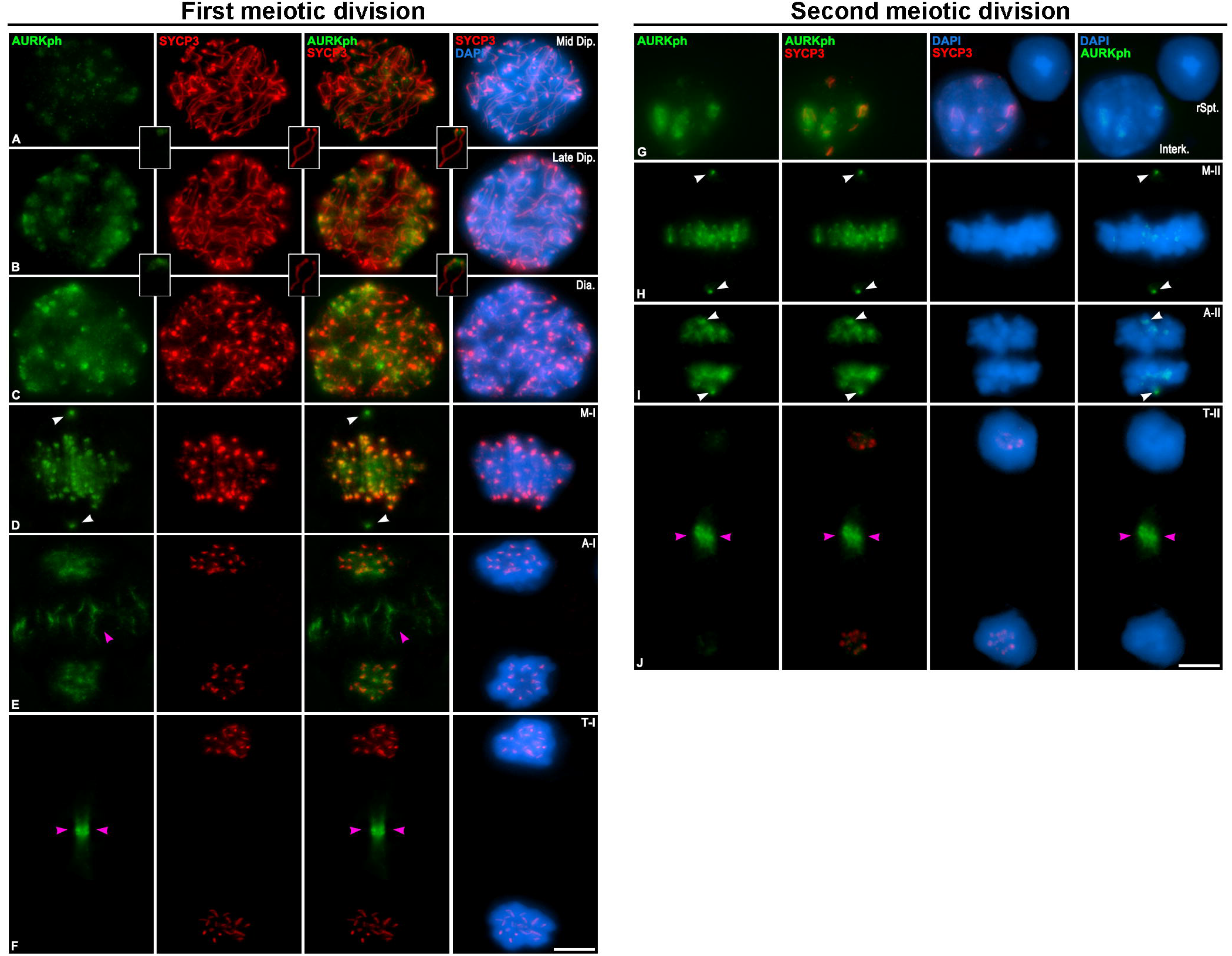
Distribution of AURKph and SYCP3 during first and second meiotic division. Double immunolabelling of AURKph (green) and SYCP3 (red) on squashed WT mouse spermatocytes on (A) mid diplotene, (B) late diplotene, (C) diakinesis, (D) metaphase-I, (E) anaphase-I, (F) telophase-I, (G) interkinesis, (H) metaphase-II, (I) anaphase-II and (J) telophase-II. Chromatin has been stained with DAPI (blue). White arrows correspond with centrosomes and magenta arrows correspond with mid body. Scale bar in J represents 10 µm.

